# Genomic occupancy of the bromodomain protein Bdf3 is dynamic during differentiation of African trypanosomes from bloodstream to procyclic forms

**DOI:** 10.1101/2022.01.11.475927

**Authors:** Ethan Ashby, Lucinda Paddock, Hannah L. Betts, Geneva Miller, Anya Porter, Lindsey M. Rollosson, Carrie Saada, Eric Tang, Serenity J. Wade, Johanna Hardin, Danae Schulz

**Affiliations:** Department of Biology, Harvey Mudd College, 301 Platt Boulevard, Claremont, CA 91711; Department of Mathematics, Pomona College, 333 N College Way, Claremont, CA 91711

## Abstract

*Trypanosoma brucei*, the causative agent of Human and Animal African trypanosomiasis, cycles between a mammalian host and a tsetse fly vector. The parasite undergoes huge changes in morphology and metabolism as it adapts to each host environment. These changes are reflected in the differing transcriptomes of parasites living in each host. While changes in the transcriptome have been well catalogued for parasites differentiating from the mammalian bloodstream to the insect stage, it remains unclear whether chromatin interacting proteins mediate transcriptomic changes during life cycle adaptation. We and others have shown that chromatin interacting bromodomain proteins localize to transcription start sites in bloodstream parasites, but whether the localization of bromodomain proteins changes as parasites differentiate from bloodstream to insect stage parasites remains unknown. To address this question, we performed Cleavage Under Target and Release Using Nuclease (CUT&RUN) timecourse experiments using a tagged version of Bromodomain Protein 3 (Bdf3) in parasites differentiating from bloodstream to insect stage forms. We found that Bdf3 occupancy at most loci increased at 3 hours following onset of differentiation and decreased thereafter. A number of sites with increased bromodomain protein occupancy lie proximal to genes known to have altered transcript levels during differentiation, such as procyclins, procyclin associated genes, and invariant surface glycoproteins. While most Bdf3 occupied sites are observed throughout differentiation, a very small number appear *de novo* as differentiation progresses. Notably, one such site lies proximal to the procyclin gene locus, which contains genes essential for remodeling surface proteins following transition to the insect stage. Overall, these studies indicate that occupancy of chromatin interacting proteins is dynamic during life cycle stage transitions, and provides the groundwork for future studies aimed at uncovering whether changes in bromodomain protein occupancy affect transcript levels of neighboring genes. Additionally, the optimization of CUT&RUN for use in *Trypanosoma brucei* may prove helpful for other researchers as an alternative to Chromatin Immunoprecipitation (ChIP).

## Introduction

The ability to adapt to different environments is vital for parasites that live in different hosts. *Trypanosoma brucei*, the protozoan parasite that causes Human and Animal African Trypanosomiasis, is one such organism. *T. brucei* lives in the bloodstream and tissues of the mammalian host (Capewell et al., 2016; Trindade et al., 2016) and multiple organs within the tsetse fly vector as it travels from the gut to the salivary gland (Roditi et al., 2016). Throughout its life cycle, the parasite adapts to each unique environment. Parasites living in the bloodstream of the mammal evade the host immune system through antigenic variation of surface proteins called Variant Surface Glycoproteins (VSGs) (Cross et al., 2014; Hovel-Miner et al., 2015). Prior to transitioning to the fly, bloodstream parasites differentiate to stumpy forms that are transcriptionally pre-adapted for making the transition to the fly gut (Rico et al., 2013). Once the parasites arrive in the midgut, they differentiate fully to procyclic forms, and VSGs on the surface are replaced by procyclin proteins (Vassella et al., 2001). Because the mammalian bloodstream and fly midgut differ in temperature, pH, and nutrient availability, the parasites undergo huge changes in morphology and metabolism. Underlying these changes are large differences in transcript levels for thousands of genes (Briggs et al., 2021; Jensen et al., 2009; Kabani et al., 2009; Nilsson et al., 2010). Researchers have made great strides in understanding how environmental signals are sensed by the parasites and what signaling pathways may be important for the transition to stumpy and procyclic forms (Cayla et al., 2020; Dean et al., 2009; McDonald et al., 2018; Mony et al., 2014; Reuner et al., 1997; Rojas et al., 2019; Vassella et al., 1997). It has also been demonstrated that RNA binding proteins play a role in differentiation processes (Jha et al., 2015; Kolev et al., 2012; Mugo and Clayton, 2017; Mugo et al., 2017; Paterou et al., 2006; Walrad et al., 2009, 2012). However, whether chromatin interacting proteins play a role in initiating or regulating changes in transcript levels necessary for transition from the bloodstream to the procyclic stage in *T. brucei* is less well understood.

Experimental observations suggest that chromatin interacting proteins might be involved in transcriptome reprogramming during the transition from bloodstream to procylic forms. Notably, inhibition of chromatin interacting bromodomain proteins in bloodstream parasites results in changes to the transcriptome that mirror those that occur as parasites transition from the bloodstream form to the procyclic form (Schulz et al., 2015). Bromodomain proteins bind to acetylated histone tails in *T. brucei* and other model organisms (Dhalluin et al., 1999; Jenuwein and Allis, 2001; Siegel et al., 2009; Yang et al., 2017) and have well established roles in gene regulation (Zaware and Zhou, 2019). Inhibition of bromodomain proteins in mammalian stem cells results in spontaneous differentiation (Di Micco et al., 2014; Horne et al., 2015). In *T. brucei*, seven bromodomain proteins (Bdfs) have been identified. Six of these proteins bind to transcription start sites at areas where polycistronic transcription units diverge (Schulz et al., 2015; Siegel et al., 2009; Staneva et al., 2021). Members of the bromodomain protein family form distinct complexes in bloodstream forms (Staneva et al., 2021). The focus of this study is Bdf3, which localizes to transcription start sites and has been shown to associate with Bdf5 and the Histone acetyltransferase HAT2 (Staneva et al., 2021). Bdf3 was chosen as a good first candidate because knockdown of this protein by RNAi results in transcriptome changes similar to those that occur during differentiation from bloodstream to procyclic forms (Schulz et al., 2015). While the localization of Bdf3 has been well characterized in bloodstream parasites, nothing is known about whether this localization is maintained during differentiation to procyclic forms. We hypothesized that Bdf3 might undergo changes in localization or occupancy at binding sites during differentiation from bloodstream to procyclic forms. Such changes in occupancy or localization could play a role in transcriptome reprogramming during differentiation.

To investigate whether localization of Bdf3 is dynamic during the transition from bloodstream to procyclic forms, we analyzed differentiating parasites using Cleavage Under Targets and Release Using Nuclease (CUT&RUN). This technique is an alternative to Chromatin Immunoprecipiation and sequencing (ChIP-seq) that avoids potential artifacts and can be performed in less time (Skene and Henikoff, 2017; Teytelman et al., 2013). Our results indicate that the CUT&RUN protocol developed in mammalian systems can be modified for successful use in *T. brucei*. We were able to use data from differentiating parasites processed by CUT&RUN to show that a *de novo* site of Bdf3 localization appears near the procyclin gene locus in differentiating parasites. More globally, occupancy at the majority of Bdf3 binding sites is transiently increased during the course of differentiation from bloodstream to procyclic forms.

## Results and Discussion

### Optimization of CUT&RUN for bloodstream form *T. brucei*

The CUT&RUN protocol was originally developed for use in mammalian cell systems (Skene and Henikoff, 2017), so we set out to adapt the protocol to *T. brucei* bloodstream parasites. In brief, the CUT&RUN protocol works as follows. Cells are permeabilized and incubated with an antibody against the protein of interest. A fusion protein of protein A and micrococcal nuclease (pA-MN) is then added. The fusion protein binds to the antibody, and when calcium is added, the fusion protein cleaves the DNA immediately surrounding the protein of interest. The small DNA fragments that are released diffuse out of the nucleus and can be collected in the supernatant following centrifugation of the permeabilized cells. These small DNA fragments are used to generate a sequencing library, which can be analyzed with statistical methods already developed for ChIP-seq. When CUT&RUN is performed using an antibody against an abundant histone protein, a characteristic ladder of bands is produced that represent 150bp increments corresponding to the size of DNA wrapped around a nucleosome (Kornberg, 1974; Skene and Henikoff, 2017). Thus, we optimized the CUT&RUN protocol in *T. brucei* using an antibody against the abundant protein histone H3.

To optimize the permeabilization step of CUT&RUN, we developed a flow cytometry assay to test permeabilization of parasite membranes. Parasites were permeabilized with various detergents and then incubated with a primary antibody against histone H3. A fluorescently labeled secondary antibody was then added and the parasites were assayed by flow cytometry. We observed anti-H3 staining following permeabilization with saponin, but did not see this staining in control samples where anti-H3 was not added (Figure 1A). Having optimized the permeabilization conditions, we next tested a series of incubation times and temperatures for the protein A micrococcal nuclease cleavage step of the protocol. We found that incubation for 5 minutes at 37ºC most efficiently produced the characteristic ladder of bands expected following successful cleavage around histone H3 (Figure 1B). However, because of the danger of non-specific cleavage at 37ºC (Skene and Henikoff, 2017), we processed experimental CUT&RUN samples for 5 minutes at 25ºC. Targeted micrococcal nuclease cleavage was dependent on the addition of calcium and did not occur when a non-specific control antibody was used (Figure 1C). These results indicate that we successfully developed a CUT&RUN protocol for use in *T. brucei* bloodstream parasites.

**Figure 1.**
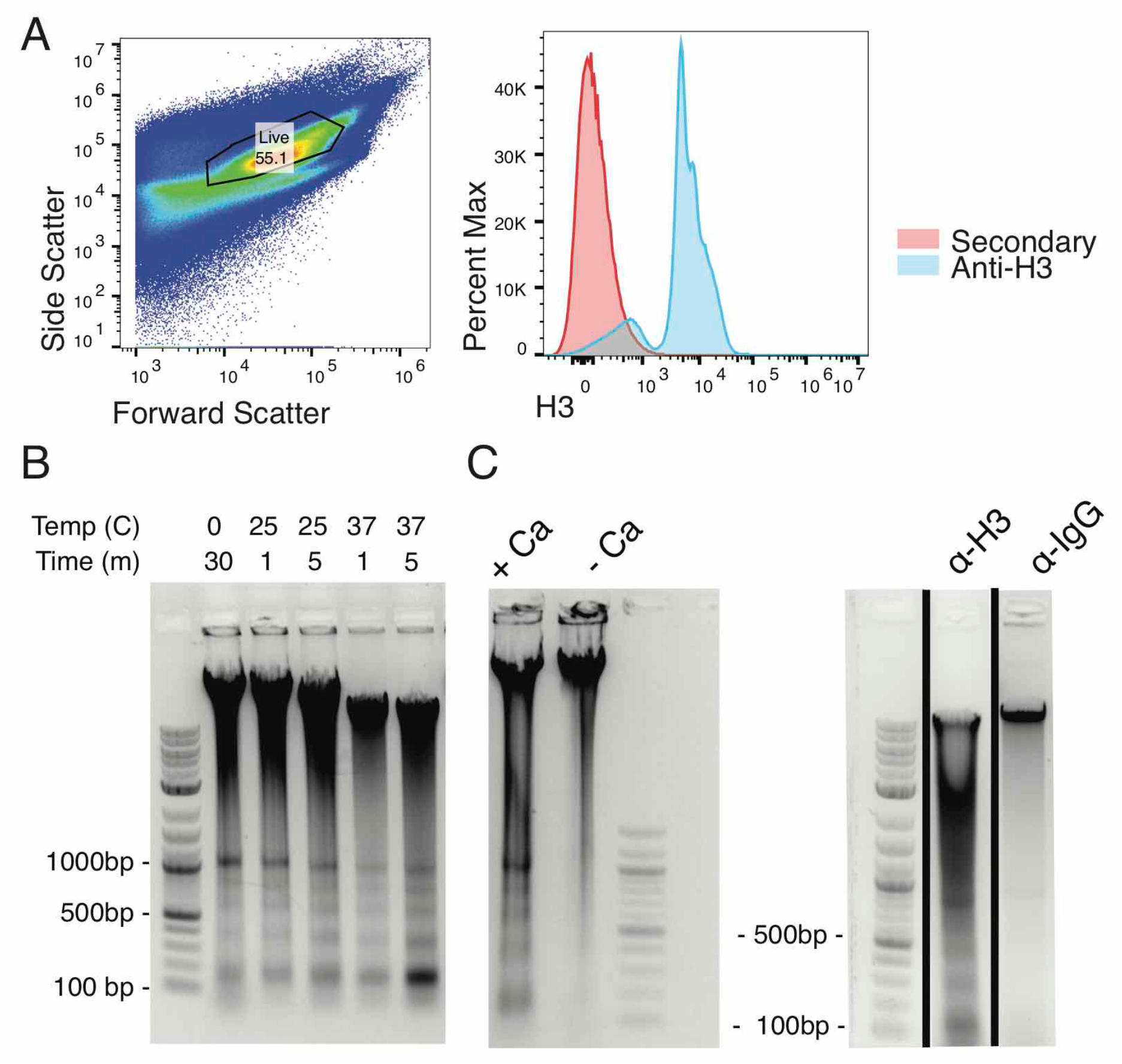
Optimization of CUT&RUN for *T. brucei*. A) Flow cytometry analysis for parasites permeabilized with 0.1% Saponin and incubated with rabbit anti-H3 primary antibody and then stained with anti-Rabbit PE. Left, forward and side scatter with live gate shown. Right, live gated parasites stained with anti-H3 (blue) or processed in parallel but with no anti-H3 added and stained with secondary antibody alone (red). B) Gel electrophoresis of DNA isolated from CUT&RUN processed parasites using an anti-H3 antibody. CaCl_2_ was added for the indicated times and temperatures to activate micrococcal nuclease cutting. C) Gel electrophoresis of DNA isolated from CUT&RUN processed parasites using an anti-H3 antibody (left panels and first lane on right panel) or an IgG antibody (right panel, second lane). Left panel shows samples processed with or without CaCl_2_ to activate micrococcal nuclease cutting. Black line indicates a gel cropped to show the indicated samples.

### Bdf3-HA tagged *T. brucei* parasites express a stumpy induction marker when grown to high density

In order to more closely approximate differentiation processes that occur in the wild, we generated EATRO1125 Antat 1.1 pleomorphic parasites with an HA-tagged allele of *Bdf3* and knocked out the remaining *Bdf3* allele (Figure 2A). Correct targeting to the endogenous *Bdf3* locus was verified using a PCR assay (Figure 2A, 2B) and our experiments were conducted with clone 6 of 24 tested clones (for simplicity, some clones are omitted from the figure). We tested our tagged parasites for their ability to generate stumpy forms by growing them to high density and measuring transcript levels of *Pad1*. We found increased transcript levels of *Pad1* when our Bdf3-HA tagged parasites were grown to high density as compared to parasites grown at low density (Figure 2C). This accords with previous work showing increased *Pad1* transcript levels in pleomorphic parasites grown to high density (Dean et al., 2009), and indicates that our Bdf3-tagged pleomorphic strain shows the expected increase in the Pad1 stumpy induction marker once stumpy formation is induced.

**Figure 2.**
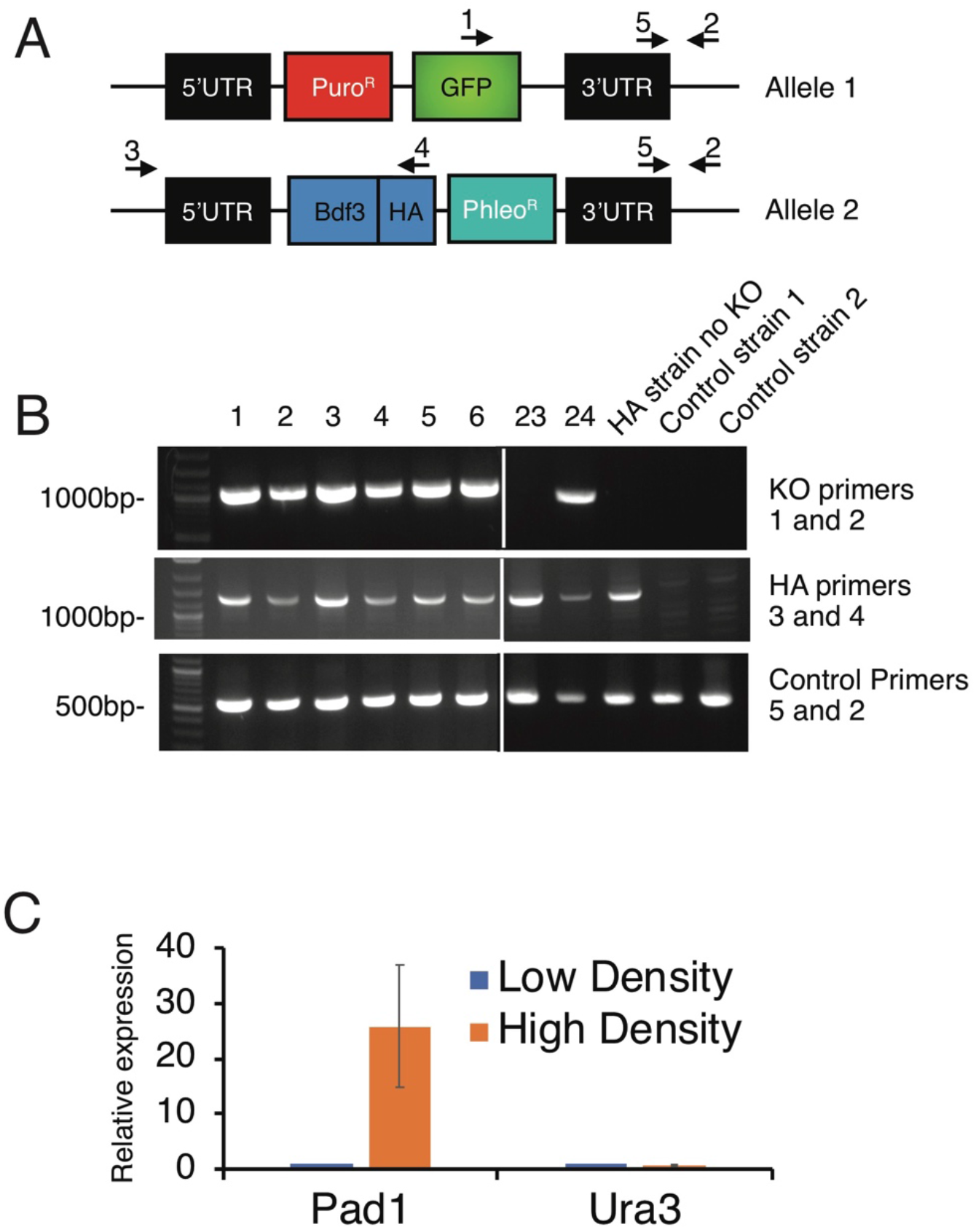
Generation of pleomorphic Bdf3-HA/Bdf3KO parasites. A) Schematic of both endogenous *Bdf3* alleles following modification with a knockout construct (top) and an HA-tagging construct (bottom). Primers used to assay for correct integration are indicated. B) Electrophoresis of genomic DNA amplified with primers to verify correct integration of the Bdf3 knockout construct (Primers 1 and 2, top panel), primers to verify correct integration of the Bdf3-HA tagging construct (Primers 3 and 4, middle panel), and control primers that should amplify all genomic DNA samples (Primers 5 and 2, bottom panel). Transformant clones are indicated as numbers. HA strain no KO indicates strain modified with the HA construct but not the KO construct. Control strains are two non-pleomorphic strains used as negative controls for the modifications. White line indicates cropping of gel photo. C) Quantitative PCR experiment to assay the transcript levels of *Pad1* and a control gene (*Ura3)* in Bdf3-HA tagged pleomorphic parasites at low density or high density.

### Bdf3 binding sites identified by CUT&RUN are similar to those identified by ChIP-seq in bloodstream parasites

Once we optimized the protocol for CUT&RUN in bloodstream parasites, we used the pleomorphic HA-tagged Bdf3 strain described above to perform CUT&RUN in bloodstream parasites using an anti-HA antibody. A non-specific IgG antibody was used as a control. We used MACS software (Zhang et al., 2008) to call peaks of Bdf3 localization using the IgG sample as a control. We compared peaks found by CUT&RUN to our published Bdf3 peaks found during ChIP-seq (Schulz et al., 2015) and found that peaks called by MACS for our CUT&RUN dataset were very similar to peaks identified in our ChIP-seq dataset (Figure 3A). 92% of peaks identified by ChIP-seq were also identified by CUT&RUN (Figure 3B) and 86% of CUT&RUN peaks were identified by ChIP-seq. Overall this suggests that CUT&RUN is a viable technique for producing localization data in *T. brucei*.

**Figure 3.**
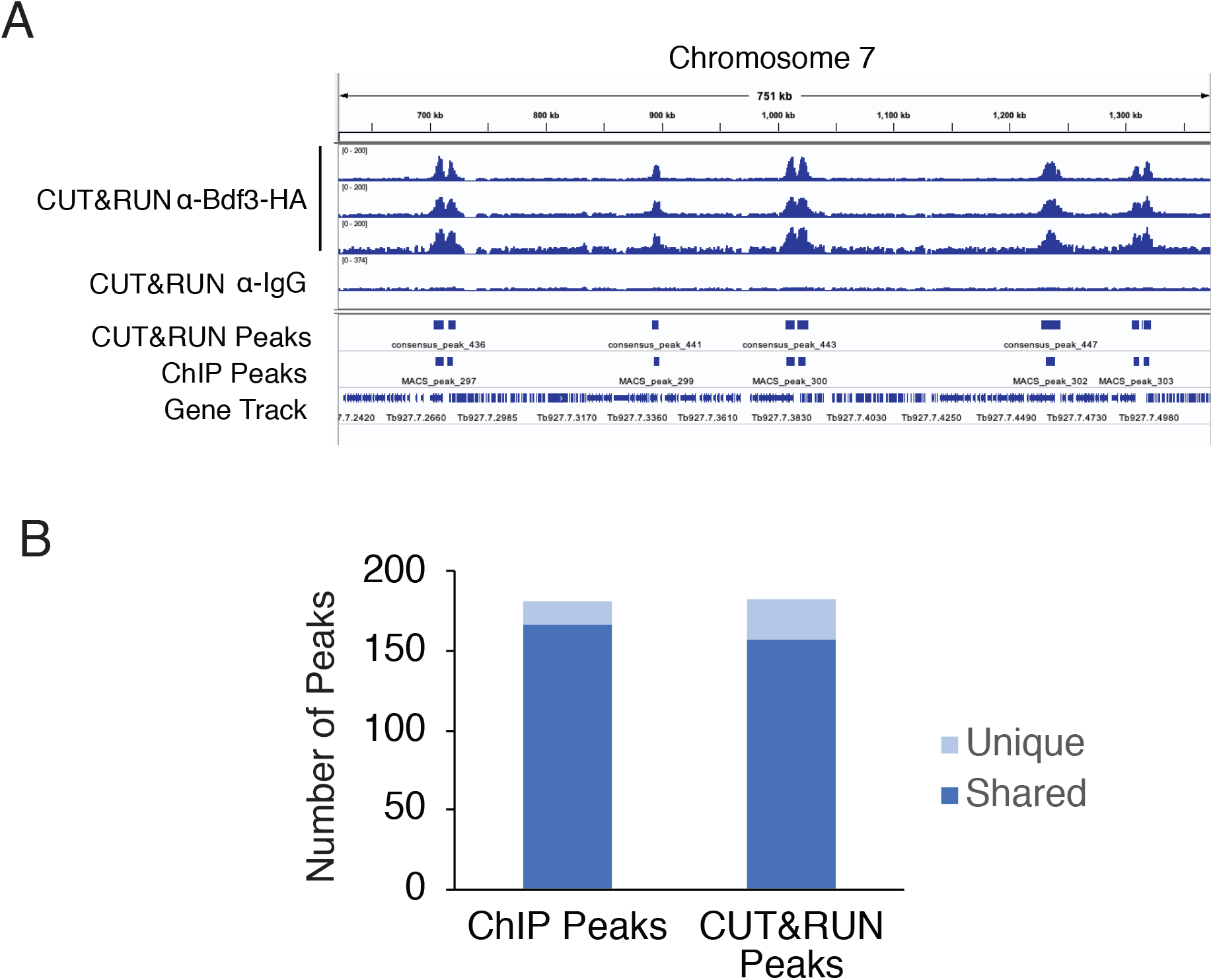
CUT&RUN identified Bdf3 binding sites are similar to those previously found by ChIP-seq. A) IGV display for a region of chromosome 7 showing sequencing tracks for three biological replicates processed for CUT&RUN using an anti-HA antibody in a pleomorphic Bdf3-HA tagged parasite line. A control sample processed with anti-IgG is also shown. Blue boxes below sequencing tracks indicate peaks of Bdf3 localization called by MACS in CUT&RUN samples and in published Bdf3-HA ChIP experiments in monomorphic strains (Schulz et al., 2015). The last row displays a gene track. B) Plot showing the number of Bdf3 ChIP-identified peaks that do or do not overlap with Bdf3 CUT&RUN-identified peaks and vice versa. Materials and methods describes details of peak merging for this analysis.

CUT&RUN may represent a more physiological state of the cell since it does not use formaldehyde crosslinking; the use of this technique could thus avoid artifacts that have been previously documented for ChIP-seq (Teytelman et al., 2013). In CUT&RUN, binding of the primary antibody to the target protein of interest occurs following permeabilization and prior to any other processing. The data produced for Bdf3 localization by CUT&RUN is quite similar to what has been seen previously for the localization of Bdf3 in bloodstream forms using ChIP-seq (Figure 3B). Bdf3 was found to localize primarily to divergent strand switch regions thought to be transcription start sites. 76% of all divergent strand switch regions (112 out of 148) were within 5kb of a Bdf3 peak. This is in accord with previous results that show Bdf3 localizing to regions where transcription is initiated (Schulz et al., 2015; Siegel et al., 2009; Staneva et al., 2021). In addition, peaks of localization called by MACS showed substantial overlap between the two techniques. This is reassuring, as it suggests that data acquired by ChIP-seq represents physiological levels of binding for the *T. brucei* proteins that have been studied using the ChIP-seq technique.

Although the data produced by ChIP-seq and CUT&RUN are similar, CUT&RUN is faster. ChIP-seq requires at least two overnight steps to immunoprecipitate complexes and reverse the formaldehyde crosslinking, with a potential third overnight step to bind primary antibodies to beads. These steps are then followed by sequencing library processing. In contrast, CUT&RUN can be performed in ∼4 hours. The development of CUT&TAG (Cleavage Under Targets and TAGmentation) has streamlined the process even further by eliminating some downstream steps associated with sequencing library preparation (Kaya-Okur et al., 2019), and has further been refined to be fully automated (Janssens et al., 2021). CUT&TAG has also been adapted for single cell chromatin studies (Wu et al., 2021), which represents an exciting future application for the technique in *T. brucei*. A recent paper used single cell sequencing to delineate transcriptome changes that occur during differentiation (Briggs et al., 2021). The use of CUT&TAG on single cells during differentiation might delineate the accompanying changes in occupancy for chromatin associated proteins throughout the differentiation process. Excitingly, CUT&RUN has been successfully adapted for use in *Toxoplasma gondii* to identify a master regulator of differentiation (Waldman et al., 2020).

### Bdf3 localizes to a region near the procyclin gene locus following differentiation

In order to ascertain whether the localization of Bdf3 is altered as parasites differentiate from the bloodstream to the procyclic form, we induced differentiation of our pleomorphic Bdf3-HA tagged strain by resuspending parasites in differentiation media, adding 6mM cis-aconitate, and incubating them at 27ºC. We harvested parasites in triplicate at 1h, 3h, 24h, and 76h post differentiation and performed CUT&RUN using an anti-HA antibody. Bloodstream parasites were also processed the same way using 5 biological replicates. Sequencing libraries were generated and the resulting reads were trimmed for quality and aligned to the *T. brucei* genome. MACS (Zhang et al., 2008) was then used to call peaks of Bdf3 localization at every time point (Figure 4, peak locations given in Supplemental Table 1). Visual inspection of the results revealed that almost all Bdf3 peaks identified in all 3 replicates of bloodstream parasites were also identified as peaks at every timepoint thereafter (Figure 4).

**Figure 4.**
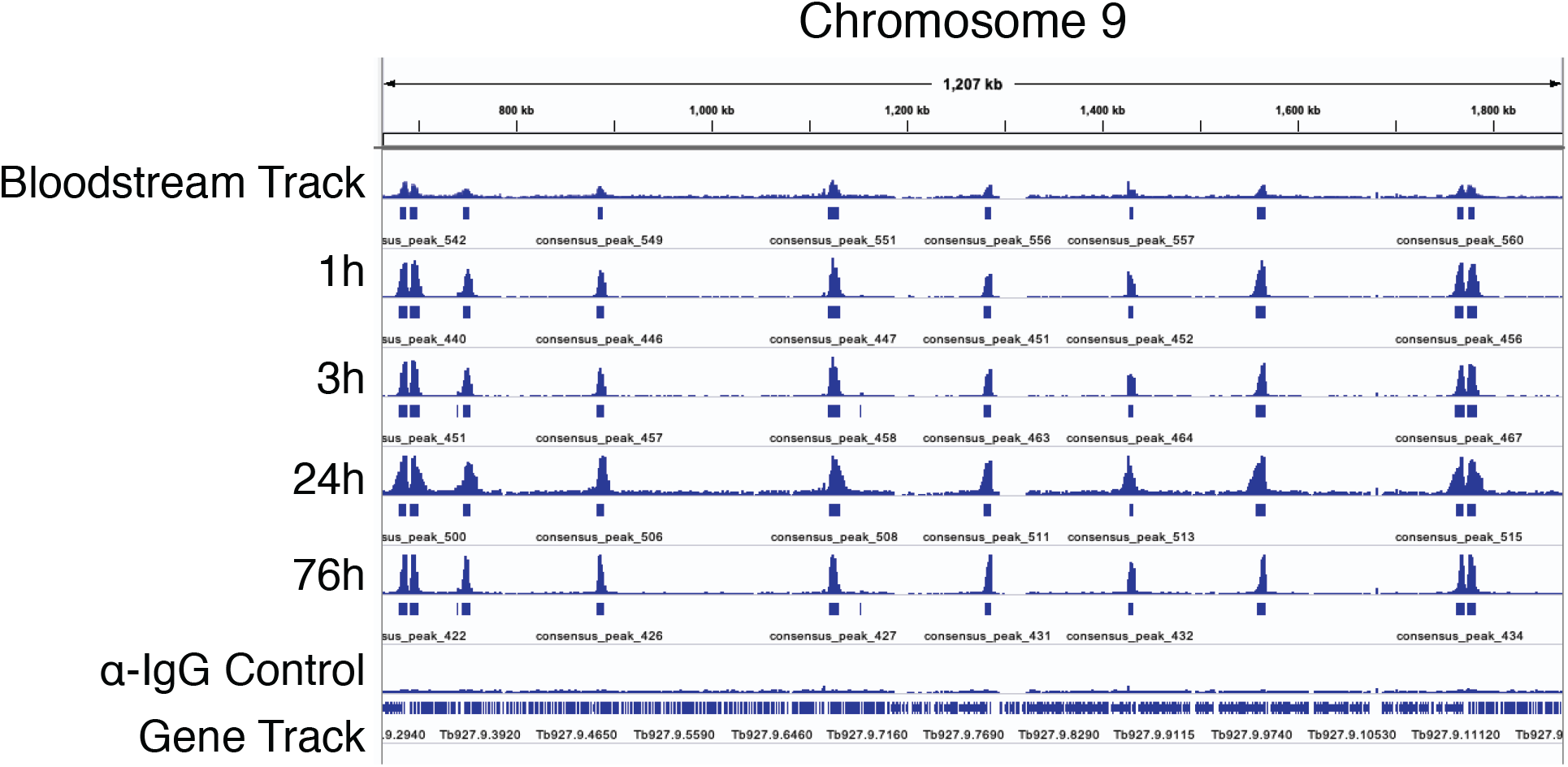
Most Bdf3 binding sites are retained throughout differentiation. IGV display for a region of chromosome 9 showing sequencing tracks for overlaid biological replicates processed for CUT&RUN using an anti-HA antibody in a pleomorphic Bdf3-HA tagged parasite line. A control sample processed with anti-IgG is also shown. Blue boxes below each sequencing track indicate peaks of Bdf3 localization identified by MACS.

While most MACS identified Bdf3 peaks were retained throughout the course of differentiation, one notable exception to this trend was found at chromosome 10 in the region of the procyclin gene locus. MACS called three Bdf3 peaks in this region at 76 hours post differentiation that were not identified as peaks earlier in the time course (Figure 5). This is especially interesting because transcripts from the *EP* and *PAG* genes near this locus are increased during the transition from bloodstream to procyclic forms so that parasites can remodel their VSG surface coat with procyclin. While RNA binding proteins have been shown to have a role in stabilizing transcripts of procyclin genes (Paterou et al., 2006; Walrad et al., 2009, 2012), the *de novo* localization of Bdf3 to this region may indicate that bromodomain proteins play a role in inducing or maintaining increased levels of procyclin transcripts during differentiation (Figure 5). This would be interesting to test in future studies, perhaps through the use of a tethering experiment where Bdf3 is artificially localized to the *EP* locus in bloodstream forms. An increase in the level of transcript for *EP1* following artificial tethering would support a model where the *de novo* appearance of Bdf3 at this locus helps to increase transcript levels of procyclin genes nearby, facilitating surface remodeling of the parasite.

**Figure 5.**
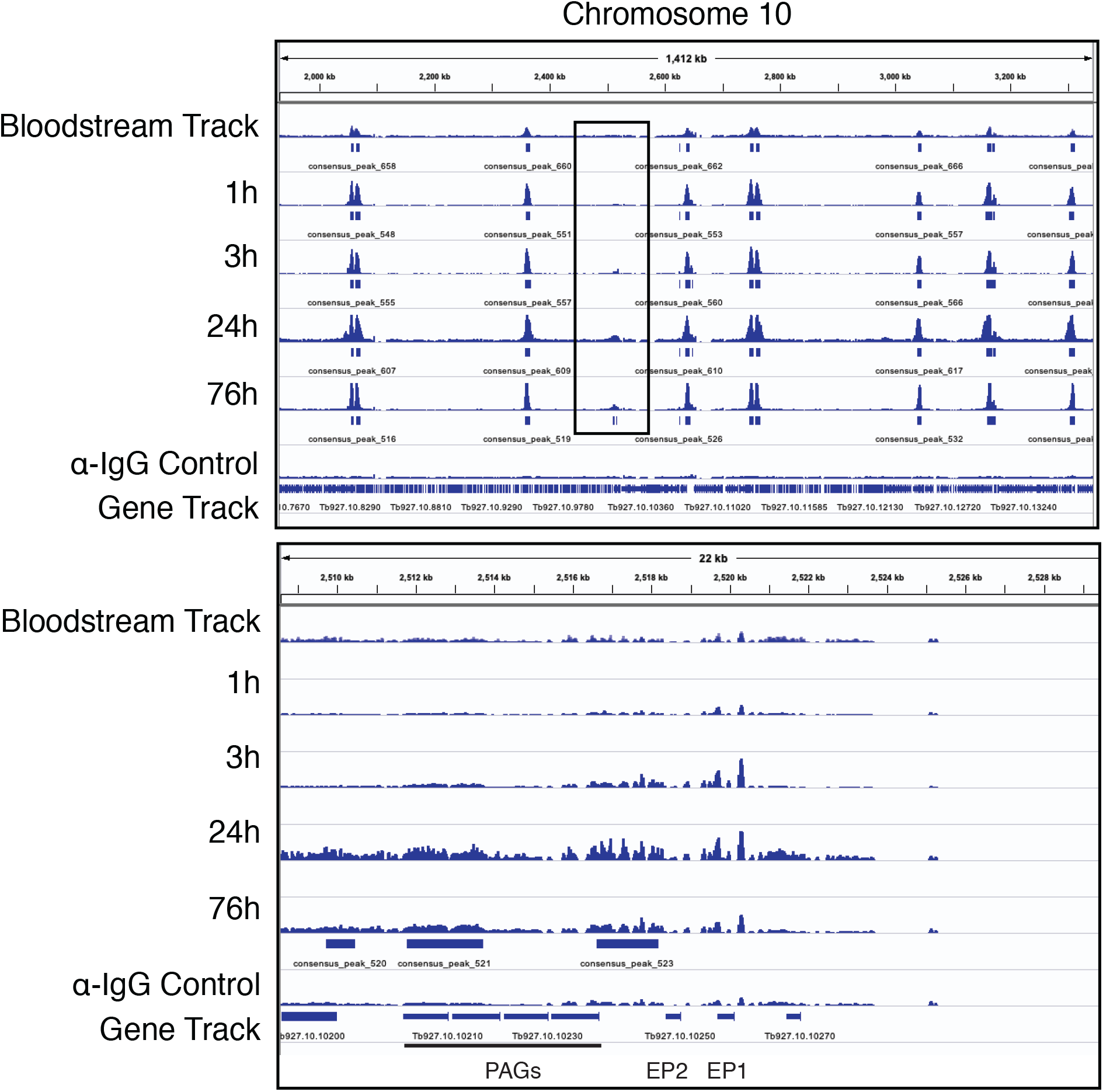
A *de novo* Bdf3 peak appears at the *EP1* locus after differentiation to the procyclic form is induced. Top panel. IGV display for a region of chromosome 10 showing sequencing tracks for overlaid biological replicates processed for CUT&RUN using an anti-HA antibody in a pleomorphic Bdf3-HA tagged parasite line. A control sample processed with anti-IgG is also shown. Blue boxes below each sequencing track indicate peaks of Bdf3 localization identified by MACS. The black square outlines the location of a new peak forming at the *EP1* locus. Bottom panel. Same as top panel except zoomed in on the *EP1* locus for chromosome 10. Three peaks are called at 76 hours that did not appear previously.

Custom scripts were used to systematically check for other instances of *de novo* Bdf3 peak formation following differentiation. While a number of sites were identified, visual inspection revealed that most of these sites were likely false positives. Many were in regions with high background in the IgG control samples. Thus, it’s likely that for some timepoints, these regions just made it over the threshold to be called by MACS as a peak, while at other timepoints they did not, leading to a false positive result. Supplemental Table 2 lists each of the sites with a numerical code given by visual inspection, with 0 indicating no evidence of *de novo* peak formation by visual inspection, 1 indicating some evidence, and 2 indicating good evidence. Sites that received a score of 1 include the *GPEET* procyclin locus on chromosome 6 and the Invariant Surface Glycoprotein (*ISG)* locus on chromosome 5. Both of the protein products for these loci are associated with differentiation processes. A similar analysis was performed to check for MACS-called Bdf3 peaks present in bloodstream forms that were not present at subsequent time points. Again, many of these loci likely represent false positives, and only two sites were scored as 1: a *VSG* locus on chromosome 5 and an *ESAG* locus on chromosome 1 (Supplemental Table 3).

### Occupancy of Bdf3 at genomic binding sites transiently increases as parasites differentiate from bloodstream to procyclic forms

Having ascertained that most sites of Bdf3 localization are retained throughout differentiation, we next wanted to measure whether occupancy at these sites is altered by quantifying tag counts within each Bdf3 peak over time. To do so, we took advantage of the DiffBind program (Ross-Innes et al., 2012; Stark, Rory, 2011) which is designed to identify changes in protein occupancy under different conditions. DiffBind takes BAM files and MACS called peaks as input, and first finds consensus peaks that are present in all biological replicates using the provided MACS files. These consensus peaks are considered regions of interest. The program normalizes the data and computes a normalized tag count for regions of interest using a 400bp region surrounding the peak summit. A region is considered to have a change in occupancy if there is a statistically significant change in tag count for the region around the peak summit at a time point after differentiation as compared to the bloodstream samples.

The method of normalization has been shown to have an outsized effect on whether a particular region is identified as having a significant change in occupancy (Stark, Rory, 2011). To circumvent this potential issue, we used four different normalization methods to identify regions with a change in occupancy over the course of differentiation. The regions identified as having a change in occupancy by all four normalization methods are considered ‘high confidence’ differentially occupied regions. The normalization methods used include (1) Reads per kilobase of transcript per million mapped reads (RPKM), (2) spike-in library size normalization, which normalizes using spiked in yeast DNA for each sample (3) background RLE, where counts are divided by sample-specific size factors determined by median ratio of gene counts relative to geometric mean per gene, a method similar to DESeq, and (4) background RLE of spiked in reads, which uses the former method with spiked-in yeast in reads. In total, we found 268 ‘high confidence’ regions that were identified as having a significant change in occupancy at a time point following differentiation when compared to the bloodstream data (Figure 6).

**Figure 6.**
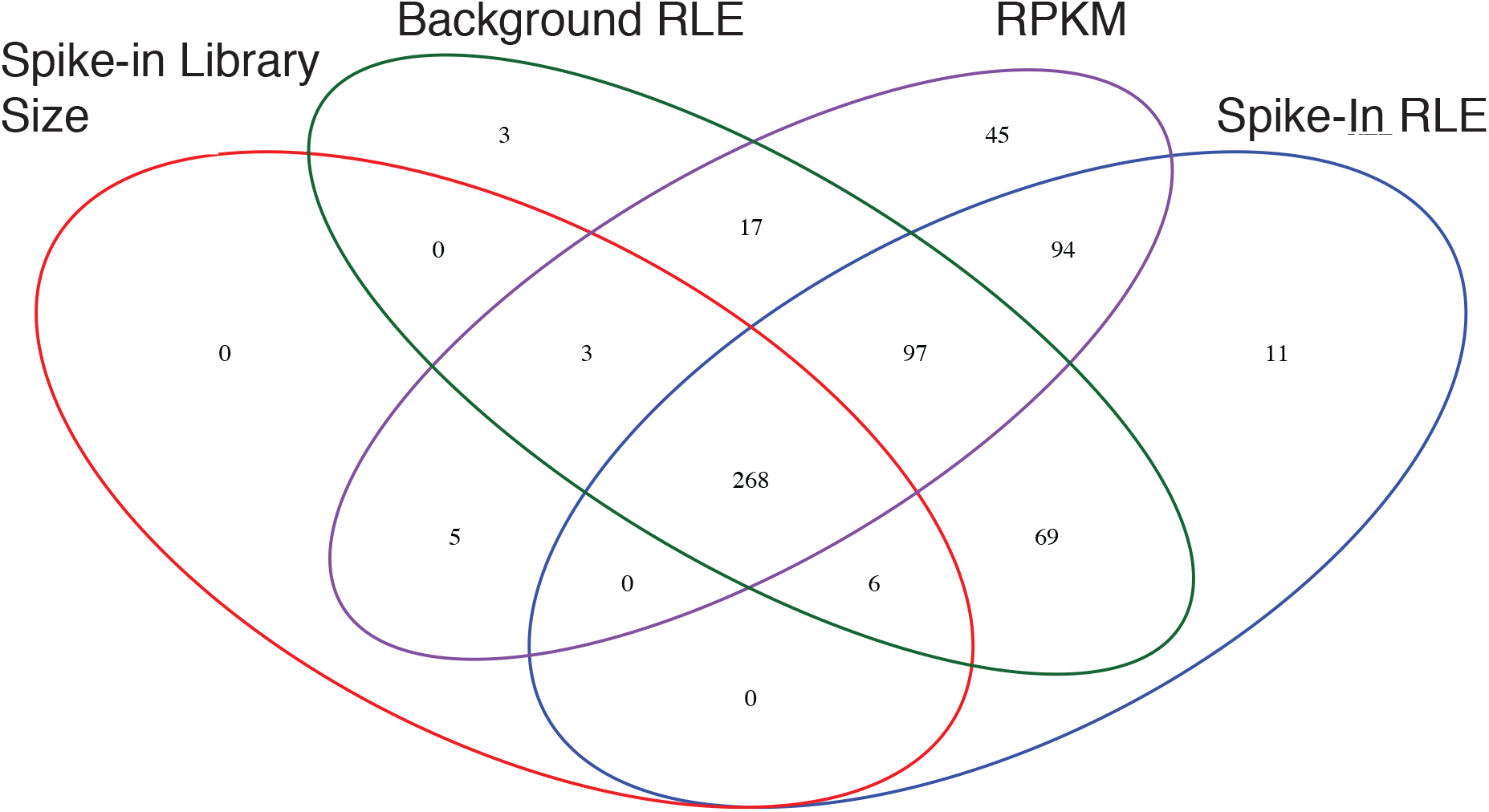
Changes in occupancy for Bdf3 are found at many loci throughout the genome following induction of differentiation from the bloodstream to the procyclic form. Venn diagram showing the number of sites identified by DiffBind as having a statistically significant change in occupancy when compared to bloodstream samples using four different normalization methods.

We plotted the normalized tag counts for each of the 268 ‘high confidence’ differentially occupied Bdf3 binding sites arrayed by chromosome (Figure 7A, Supplemental Figure 1, Supplemental Table 4). There was a remarkable similarity to the pattern of occupancy for these differentially bound regions over time, with most regions showing a peak in occupancy at 3 hours post differentiation and a decrease in occupancy thereafter. In order to ensure that the peak in occupancy observed for Bdf3 at 3 hours post differentiation was not an artifact, we randomly shuffled Bdf3 binding site locations using bedtools. The regions were shuffled such that each shuffled site was retained on the same chromosome as the original site and all regions identified as consensus sites were excluded. These shuffled regions were then run through the DiffBind program and normalized the same way as the Bdf3 consensus sites (Figure 7B, Supplemental Figure 2, Supplemental Table 5). When tag counts for shuffled control regions were plotted over time, we did not observe the same peak in the number of tag counts at the 3-hour time point, indicating that the peak in Bdf3 occupancy that occurs 3 hours after differentiation represents a genuine change in the amount of Bdf3 found at that location, either at the single cell or the population level.

**Figure 7.**
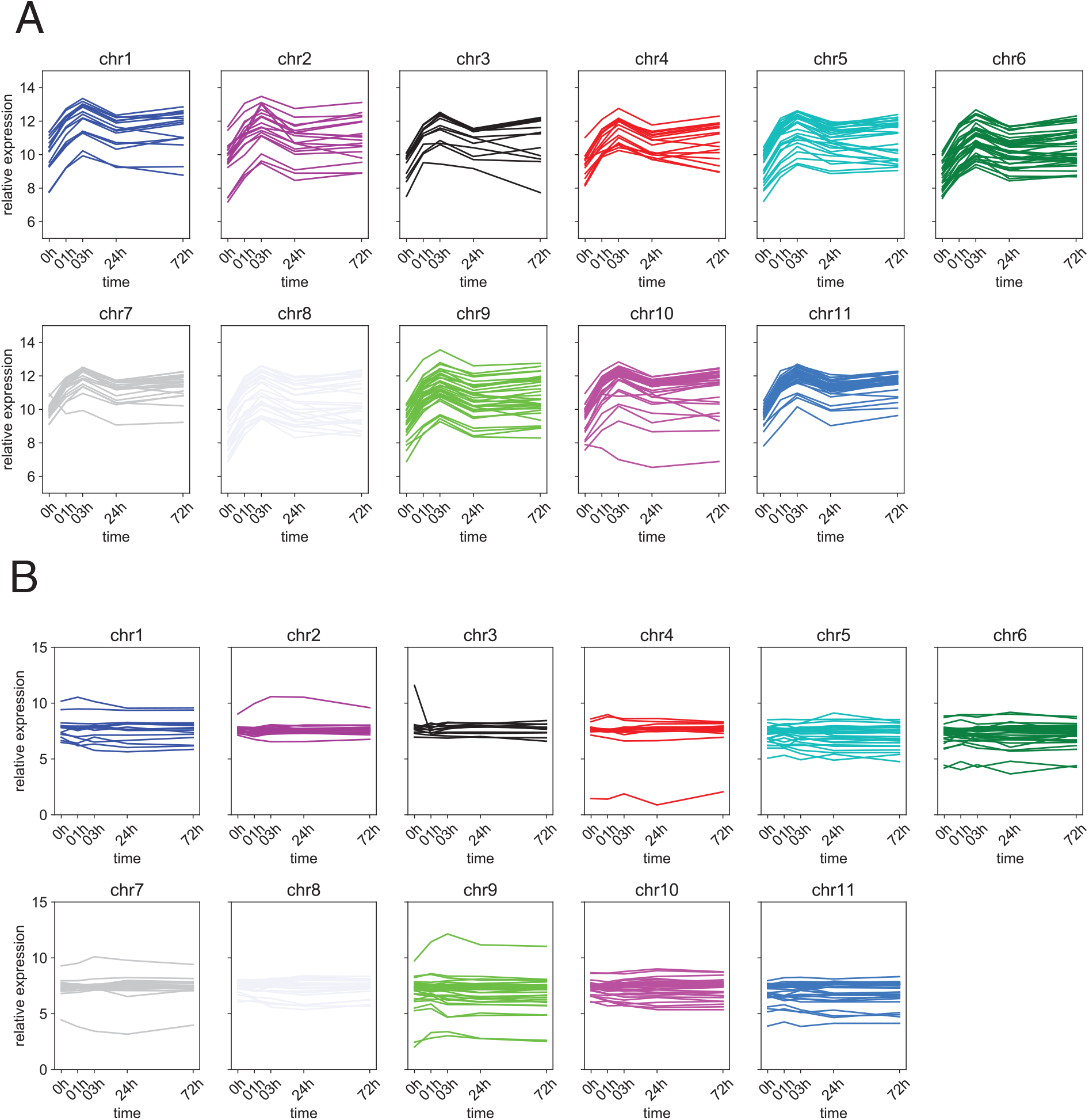
Bdf3 occupancy peaks at 3 hours after induction of differentiation. A) Plots of log_2_ normalized tag counts over time for each ‘high confidence’ Bdf3 binding site with a statistically significant change in occupancy after induction of differentiation to the procyclic form using background RLE normalization and arrayed by chromosome. B) Plots of log_2_ normalized tag counts over time for control regions that are not identified as Bdf3 binding sites using background RLE normalization.

The fact that the peak of Bdf3 occupancy is reached 3h following addition of cis-aconitate is especially interesting in light of work showing that commitment to differentiation occurs 2-3h hours after cis-aconitate treatment (Domingo-Sananes et al., 2015). This group further demonstrated that protein synthesis is required to generate ‘memory’ of exposure to the differentiation signal. One possible model is that bromodomain protein occupancy at transcription start sites facilitates commitment at 3h following differentiation by an unknown mechanism. For instance, if increased occupancy led to an increase in transcript levels, this could aid in the protein synthesis required for commitment to differentiation.

Of the 268 ‘high confidence’ sites with changes in Bdf3 occupancy, 162 are within 5kb of a divergent strand switch region considered to be sites of transcription initiation in *T. brucei*. This might be an underestimate of the true number of dynamically occupied Bdf3 sites at divergent strand switch regions because we used only unique alignments, and thus we lack data for highly repetitive regions with divergent strand switch regions near the ends of the chromosomes. Of 148 divergent strand switch regions we examined, 60% were within 5kb of a ‘high confidence’ dynamically occupied Bdf3 site. The effect of the increased occupancy at sites of Bdf3 localization is not yet known for *T. brucei*, but in other systems, changes in occupancy for chromatin binding proteins can reflect an underlying change in particular histone tail modifications that are bound by the chromatin binding protein (Ross-Innes et al., 2012). In our particular case, a change in histone acetylation at transcription start sites could result in the observed increase in Bdf3 occupancy at these sites as parasites begin differentiation. Future studies could investigate the acetylation levels at transcription start sites after the induction of differentiation from the bloodstream to the procyclic form. A transient increase in acetylation at transcription start sites might result in a corresponding transient increase in bromodomain protein localization at these sites that is observed for Bdf3 (Figure 7). This leaves us with the question of how an increase in acetylation might support the differentiation process. One model is that such an increase in acetylation might increase genomic transcription overall, and thus help the parasites exit the relatively quiescent cell cycle arrested stumpy stage and transition to the dividing procyclic stage. In other systems, quiescence results in a global decrease in acetylation levels (McKnight et al., 2015; Swygert et al., 2019; Young et al., 2017). Specific chromatin remodeling enzymes are required to generate hypertranscription necessary for exiting the quiescent state (Cucinotta et al., 2021). If an increase in acetylation levels and increased transcription are necessary during the transition from the quiescent stumpy form to the cycling procyclic form, this could result in the observed increase in Bdf3 occupancy following the differentiation cue. Bromodomain protein mediated regulation of global Pol II transcript levels has been demonstrated in *Leishmania* (Jones et al., 2021). RNA-seq analyses are typically normalized in such a way as to obscure global increases or decreases in transcript levels (Evans et al., 2018). Thus, it might be interesting to examine whether the exit from the stumpy to the procyclic stage results in a global increase in transcript levels for Pol II regulated genes.

Bromodomain inhibition in bloodstream parasites using the small molecule inhibitor I-BET151 results in changes in the transcriptome that mirror those that occur during differentiation from the bloodstream to the procyclic stage in many ways, including an increase in *EP1* transcript levels (Schulz et al., 2015). We observed a sharp decrease in bromodomain protein occupancy for at least 268 Bdf3 binding sites in the period between 3h and 24h after the initiation of differentiation (Figure 7). One model for why bromodomain inhibition may trigger transcriptome changes akin to differentiation is that binding of the drug to its bromodomain protein target causes a sharp decrease in occupancy for bromodomain proteins in bloodstream forms as is seen at 3 hours post differentiation (Figure 7). We did observe decreased enrichment of Bdf3 in I-BET151 treated bloodstream parasites at several sites, consistent with this model (Schulz et al., 2015). If the decrease in Bdf3 occupancy in differentiating parasites is partly responsible for promoting the transition to a procyclic-specific transcriptome, then an artificial I-BET151 induced decrease in Bdf3 occupancy might also produce the observed changes in transcript levels for procyclic-associated genes.

A number of genes known to be associated with differentiation were within 5kb of a dynamically occupied Bdf3 site (Supplemental Table 6 lists all genes within 5kb of ‘high confidence’ Bdf3 sites and separates out those identified by Queiroz et al. as having altered transcript levels during differentiation (Queiroz et al., 2009)). These genes include, but are not limited to, invariant surface glycoproteins (ISGs), procyclin associated genes (PAGs), the procyclin GPEET and EP3 genes, Expression Site Associated Genes (ESAGs), Variant Surface Glycoproteins (VSGs), COX genes, flagellar genes, and adenylate cyclases (Supplemental Table 6).

In conclusion, we have adapted the CUT&RUN technique for use in *T. brucei* parasites, and used it to track *de novo* Bdf3 peak formation and changes in occupancy at Bdf3 binding sites during the transition from the bloodstream to the procyclic form. The mechanistic details for how changes in bromodomain protein occupancy might promote differentiation is an exciting area for future study.

## Materials and Methods

### Parasite Culture and Strain Generation

Bloodstream parasites were cultured in HMI-9 at 37ºC with 5% CO_2_. Differentiation was induced by resuspending parasites in Differentiation Media (DTM) (Ziegelbauer et al., 1990), adding 6mM cis-aconitate, and incubating parasites at 27ºC. The Bdf3-HA/Bdf3 KO strain was generated from EATRO 1125 AnTat1.1 90:13 (Engstler and Boshart, 2004). The pMOTAG5H Bdf3-HA construct (Schulz et al., 2015) was linearized and introduced into parasites using an AMAXA nucleofector kit. Correct integration was verified using PCR with primers: 1) HA rev (tatgggtacgcgtaatcaggcaca) and 2) upstream Bdf3 5’UTR for (tgttgcaggatattgtgagtga). After this the pyrFEKO Puro/GFP Bdf3 KO construct was linearized and transfected into Bdf3-HA tagged parasites as above and verified with primers: 1) GFP for (ctacaacagccacaaggtctat) and 2) downstream Bdf3 3’ UTR rev (aaaccgcaaagtgatgaatgg). Control primers shown in figure are 1) Bdf3 3’ UTR for (cttgtagacagcggcatggttgg) and 2) downstream Bdf3 3’ UTR rev (aaaccgcaaagtgatgaatgg).

### CUT&RUN

All spins prior to permeabilization were performed at 10ºC and 2738g. Spins after the permeabilization step occurred at 10ºC and 4602g. 50-75 million parasites were harvested via centrifugation at 10ºC and washed in 1ml NP buffer containing 0.5mM spermidine, 50mM NaCl, 10mM Tris-HCl pH 7.5, and protease inhibitors. Parasites were spun and permeabilized using 100μl NP buffer supplemented with 0.1% saponin (vol/vol) and 2mM EDTA. 5μg of anti-HA (Sigma H6908) or anti-IgG (Fisher 02-6102) control antibody was added and samples were rotated for 45m at 25ºC. For histone experiments, 1.5μl rabbit anti-H3 (a kind gift from Christian Janzen) was added. Samples were washed and pelleted twice with 1ml NP buffer. A volume corresponding to 1.4% of the sample was removed for flow cytometry analysis. Following the second wash, samples were resuspended in 100μl NP buffer and 0.5μl proteinA-Micrococcal nuclease was added (a kind gift from Steven Henikoff). Samples were rotated for 5m at 25ºC. Samples were washed twice in 1ml NP buffer as above and resuspended in 100μl NP buffer. CaCl_2_ was added to a final concentration of 2mM and nuclease digestion occurred for 5m at 25ºC. 100μl of 2X STOP buffer (20mM EDTA, 20mM EGTA) with yeast spike-in DNA was immediately added. Samples were incubated for 10m at 37ºC to release insoluble nuclear chromatin. Samples were pelleted and the supernatant containing the DNA was saved. SDS was added to a final concentration of 0.1%, proteinase K was added at 165μg/ml, and RNaseA was added to 6.5μg/ml. Samples were incubated at 70ºC for 10m and purified using phenol chloroform or Ampure XP beads at 1.8X according to the manufacturer’s instructions.

### Flow Cytometry

All flow cytometry was performed on a Novocyte 2000R from Acea Biosciences. Parasites were resuspended in 100μl HMI-9 and stained for 10 minutes on ice with mouse anti-rabbit IgG PE (Santa Cruz sc-3753). Cells were washed twice in HMI-9 prior to analysis.

### Quantitative PCR Analysis

To quantify transcript levels of *PAD1*, parasites were grown to a density of 190,000 cells/ml for low density samples or >= 1 million cells/ml for high density samples. RNA was extracted from low or high density parasite populations using RNA Stat-60 (Tel-Test) following the manufacturer’s protocol and quantified on a NanoDrop2000c. 2.5μg of RNA was used to generate cDNA using the SuperScript IV VILO Master Mix (Fisher Scientific 11756050) according to the manufacturer’s protocol. cDNA was amplified using 2X Sybr green master mix (Life Technologies 4309155) and primers and quantified on an Eppendorf Realplex2 instrument. Primers used were *PAD1*: gaccaaaggaaccttcttcct and cactggctcccctaagct, *URA3*: cggcagcagttctcgagt and tggcgtgtaccttgaggc.

### Generation of Sequencing Libraries

Sequencing libraries were generated using the NEBNext Ultra II DNA Library Prep Kit for Illumina (E7645) according to the manufacturer’s instructions with the following modification: Samples were incubated with USER enzyme immediately prior to PCR, rather than at an earlier step. NEBNext Multiplex Oligos for Illumina were used to prepare multiplex samples (e.g. E7710).

### CUT&RUN Sequencing Analysis

Sequencing was performed at the UCLA Technology Center for Genomics and Bioinformatics using an Illumina HISEQ 3000 with 50bp single end reads. CUT&RUN fastq files were trimmed using TrimGalore (http://www.bioinformatics.babraham.ac.uk/projects/trim_galore/) and aligned to the Tb927v5.1 genome using bowtie (Langmead et al., 2009) and requiring unique alignments using the following command: bowtie --best --strata -t -v 2 -a -m 1. Spike-in reads were aligned to the yeast sacCer3/R64 genome. MACS (Zhang et al., 2008) was used in broad peak mode to identify peaks of Bdf3 localization using an IgG control with the following arguments -g 23650671, --keep-dup all, --nomodel, and –broad. The DiffBind (Stark, Rory, 2011) package was used to identify regions with a change in Bdf3 occupancy. The GreyList ChIP package eliminated problematic regions from the analysis (0.66% of the genome). Four different normalization methods were then used to obtain normalized read counts for areas of interest determined by MACS: Spike-in library size, Spike-in RLE, Background RLE (not using spike-in reads), and RPKM. Both RLE methods adjust for regional ‘background’ read frequency by counting reads in non-overlapping 15kb genomic bins. This approach adjusts for broad patterns in background read enrichment while avoiding spurious adjustment to locally enriched regions. DiffBind was then used to identify regions with a statistically significant change in read counts (occupancy) at each time point using bloodstream values as a control with a cutoff of p_adj_ < 0.05 using the Benjamini-Hochberg adjustment to control the FDR. Once these regions were identified using each normalization method, a Venn diagram was used to identify ‘high confidence’ regions with changes in occupancy that were identified using all four normalization methods. To generate control regions, we randomly shuffled ‘high confidence’ regions for each chromosome using bedtools (Quinlan and Hall, 2010). Shuffled regions were maintained on the same chromosome and peaks of Bdf3 localization identified by MACS were excluded. Control regions were run through the same DiffBind pipeline to obtain normalized read counts, which were plotted for each control region.

Overlap between ChIP and CUT&RUN-identified peaks was performed by first merging peaks that were within 5kb in each dataset and then identifying unique vs overlapping peaks using bedtools (Quinlan and Hall, 2010).

*De novo* peak formation was ascertained using a custom script. The program took as input the set of MACS called Bdf3 consensus peaks at each time point and identified sites that were not called as Bdf3 peaks in bloodstream forms but subsequently were called as peaks at 2 consecutive timepoints following differentiation. The script for ‘disappearing’ peaks took the same input and identified sites that were called as peaks in bloodstream parasites and not called as peaks at two consecutive time points following differentiation.

## Acknowledgements

We would like to thank Christian Janzen for his kind gift of the anti-H3 antibody. We also thank Monica Mugnier for helpful comments on the manuscript.

---

**Supplemental Figure 1**. Plots of log_2_ normalized tag counts over time for each ‘high confidence’ Bdf3 binding site with a statistically significant change in occupancy after induction of differentiation to the procyclic form using indicated normalization.

**Supplemental Figure 2**. Plots of log_2_ normalized tag counts over time for control regions that are not identified as Bdf3 binding sites using background indicated normalization.

**Supplemental Table 1**. Normalized tag counts over time for Bdf3 peaks identified by MACS.

**Supplemental Table 2**. Analysis of Bdf3 binding sites that appear *de novo* during differentiation.

**Supplemental Table 3**. Analysis of Bdf3 binding sites that are present in the bloodstream form and not called by MACS following differentiation.

**Supplemental Table 4**. Normalized tag counts over time for Bdf3 sites with dynamic occupancy identified by DiffBind using all four normalization methods.

**Supplemental Table 5**. Normalized tag counts over time for control regions that are not Bdf3 binding sites.

**Supplemental Table 6**. Genes within 5kb of a ‘high confidence’ dynamically occupied Bdf3 site.

