## Supplemental Figures for "Genomic occupancy of the bromodomain protein Bdf3 is dynamic during differentiation of African trypanosomes from bloodstream to procyclic forms"

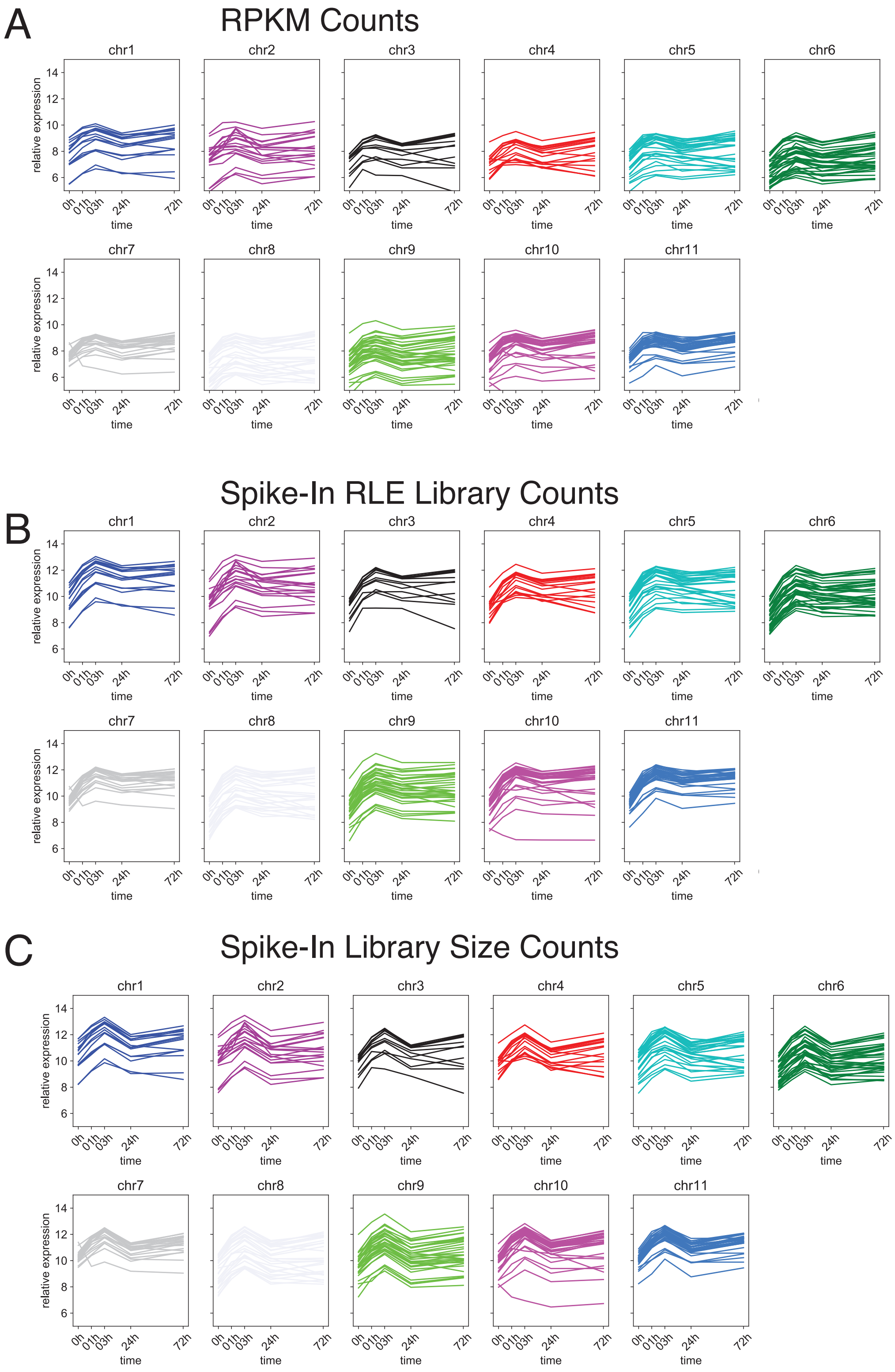


**Supplemental Figure 1.** Plots of $log_2$ normalized tag counts over time for each ‘high confidence’ Bdf3 binding site with a statistically significant change in occupancy after induction of differentiation to the procyclic form using indicated normalization.

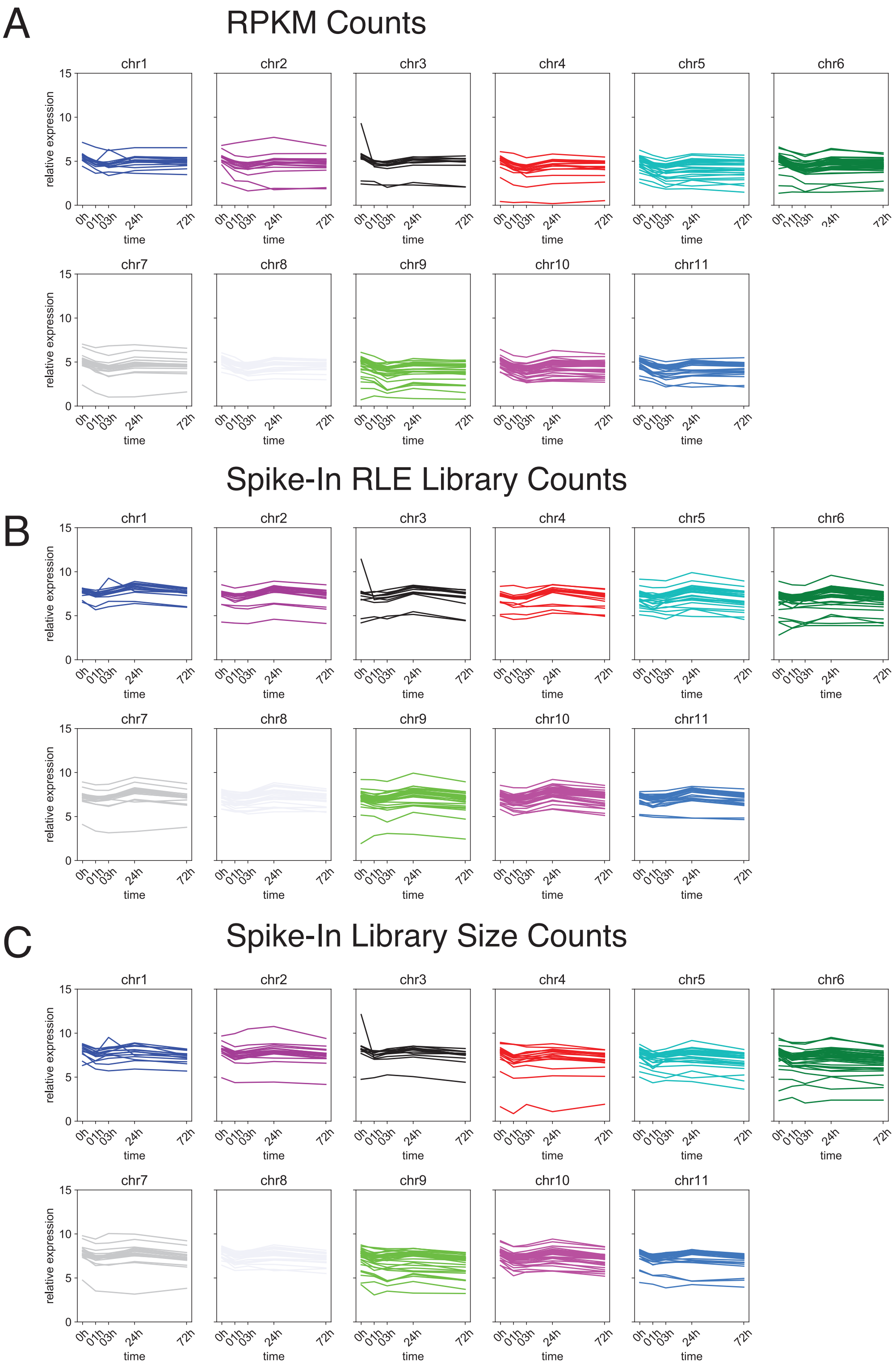


**Supplemental Figure 2.** Plots of $log_2$ normalized tag counts over time for control regions that are not identified as Bdf3 binding sites using background indicated normalization.
